# S4, a Selective Androgen Receptor Modulator, Suppresses Breast Cancer Progression via Cell Cycle Arrest, Apoptosis, and Metabolic Alterations

**DOI:** 10.1101/2025.05.05.652263

**Authors:** Mervenur Yavuz, Turan Demircan

## Abstract

Breast cancer (BC) remains a leading cause of cancer-related mortality worldwide, highlighting the need for novel therapeutic strategies. Androgen receptor (AR) signaling has been implicated in BC progression, making it a potential target for treatment. Selective androgen receptor modulators (SARMs) have gained attention as alternatives to traditional hormone therapies. However, the effects of S4, a SARM, on BC have not been explored. This study investigates the impact of S4 on BC cell viability, proliferation, clonogenicity, migration, apoptosis, and cell cycle progression in MCF-7 and MDA-MB-231 cells. The results demonstrate that S4 significantly reduces BC cell viability in a dose-dependent manner, with IC_50_ values of 0.094 mM (MCF-7) and 0.067 mM (MDA-MB-231) after 24 hours. S4 suppresses clonogenicity and migration while promoting apoptosis and cell cycle arrest, with MCF-7 cells exhibiting S-phase arrest and MDA-MB-231 cells G0/G1-S arrest. Gene expression analysis reveals upregulation of tumor suppressor genes (*CDKN1B* and *PUMA* in MCF-7 cells, *TP53, CDKN1A*, and *BAX* in MDA-MB-231 cells, and *GADD45A* in both cells) and downregulation of oncogenes (*ANKRD1, EDN1, CCND1* in MCF-7 cells, *CXCL2* in MDA-MB-231 cells, *CDK-6* and *ATM* in both cells), supporting the anti-carcinogenic effects of S4. Metabolomics profiling highlights significant alterations in phosphatidylcholine biosynthesis, catecholamine biosynthesis, and nucleotide metabolism, indicating a metabolic shift upon S4 treatment. These findings provide the first evidence of S4’s potential as an anti-cancer agent in BC, suggesting that it exerts its effects by modulating gene expression and cellular metabolism. Further *in-vivo* studies are warranted to validate its therapeutic potential.

## Introduction

Breast cancer (BC) is one of the most fatal cancers around the world and among women which still lacks the effective and beneficial treatment options [1]. The molecular heterogenicity of the BC has evolved the standard therapeutic options to more biologically directed therapies to prolong survival and reduce the tumor burden. Despite the improvements in BC therapy, recent research still emphasizes the necessity of the novel drugs and/or multidisciplinary management for its therapy [2].

The androgen receptor (AR) is a steroidal nuclear receptor type specifically plays significant roles in male sexual development, bone density, muscle mass, strength, and cognition [3]. In a ligand-dependent manner, the activation of AR through androgens results in modulation in DNA transcription of androgen-responsive elements within target genes which leads to the regulation of gene transcription responsible for differentiation, proliferation, apoptosis, or angiogenesis [4]. Besides, AR could be activated through the crosstalk with key signaling pathways including PI3K/Akt, ERK, and mTOR in a ligand-independent manner [5]. In addition to its function in male sexual development, AR also has essential roles in normal breast physiology and BC pathology. More than 70% of the BC was previously found to be express AR and therefore, in different BC subtypes AR was considered as a prognostic value [6].

Although AR and androgens are vital for the normal development of various tissue including breast, it is also evidenced that they are also responsible for stimulating tumorigenesis in prostate, heart and liver [7]. These pathology-inducing roles of androgens lead to the development of tissue-selective non-steroidal AR modulating agents, namely selective androgen receptor modulators (SARMs), to provide therapeutic benefits of androgen without off-target activities and androgenic side effects such as prostate enlargement, acne, and virilization [8]. In addition, SARMs have the ability to extend the androgen therapy to women since androgens could aromatize into estrogen which could cause undesirable side effects [9]. Moreover, we previously examined the effects of S4 on various cancer types and demonstrated its anti-carcinogenic potential [10–13].

In this study, we investigated the effects of S4 on BC cell viability, proliferation, clonogenicity, migration, apoptosis, and cell cycle regulation in both Estrogen receptor (ER)+/AR+ (MCF-7) and triple-negative (MDA-MB-231) breast cancer models. Additionally, we performed gene expression and metabolomics analysis to determine the molecular alterations induced by S4 treatment. Our results suggest that S4 suppresses BC carcinogenicity by reshaping gene expression and inducing metabolic reprogramming. These novel findings underscore the anti-cancer potential of S4 in BC cells and lays the groundwork for further investigation of SARMs as promising therapeutic agents in breast cancer treatment.

## Methods & Materials Cell culture maintenance

The human breast adenocarcinoma cell lines MCF-7 (cat. no. HTB-22, ATCC) and MDA-MB-231 (cat. no. HTB-26, ATCC) cells were maintained in Dulbecco’s Modified Eagle’s Medium (DMEM) (cat. no. D6429, Sigma-Aldrich), supplemented with 10% fetal bovine serum (FBS) (cat. no. A4736401, Thermo Fisher Scientific) and 1% penicillin/streptomycin (cat. no. 15140122, Thermo Fisher Scientific). The cells were incubated at 37°C in a humidified incubator with 5% CO_2_, and passaging was performed upon reaching 80% confluency. Cell proliferation was routinely monitored using an inverted microscope. S4 (S)-3-(4-Acetylaminophenoxy)-2-hydroxy-2-methyl-N-(4-nitro-3-trifluoromethylphenyl) propionamide, (cat. no. 78986, Sigma-Aldrich) was dissolved in dimethyl sulfoxide (DMSO) to prepare stock solutions, which were stored at -80°C for up to six months and diluted in culture medium before use.

### Cell viability assay

Cell viability was assessed using the CellTiter 96® Non-Radioactive Cell Proliferation Assay Kit (cat. no. G4000, Promega). BC cells were seeded into 96-well plates at a density of 0.1 × 10^5^ cells per well in 100 μL of culture medium and incubated for 24 hours. Subsequently, the medium was replaced with various concentrations of S4 (0.0001 mM to 0.4 mM), and cells were incubated for an additional 24 hours. Negative controls consisted of culture medium without drug, while positive controls contained 10% DMSO. The dye solution was added to each well, and after 4 hours, the solubilization solution/stop mix was introduced to halt the reaction. Absorbance was recorded at 570 nm. The experiment was repeated under the same conditions with a cell density of 0.05 × 10^5^ cells for 48 hours. IC_50_ values were determined by fitting dose-response curves using the “drc” package in R programming language, as previously described [11].

### Colony formation assay

To evaluate the colony-forming potential of cells post-drug treatment, a colony formation assay (CFA) was conducted. Cells (2 × 10^3^ per well) were plated in 96-well plates containing 100 μL of culture medium. After 24 hours, the medium was replaced and replenished every two days until the cells reached 80% confluency or higher. Colonies were fixed with 100% methanol for 20 minutes at room temperature, stained with 0.2% Crystal Violet for 15 minutes, and washed twice with ddH_2_O to remove excess stain. The plates were air-dried, and digital images were acquired using a bright-field microscope (SOPTOP ICX41). The ColonyArea plugin in ImageJ software was utilized for quantification of colony number and intensity.

### Soft agar assay

A soft agar assay was performed to examine the anchorage-independent growth of BC cells. A 5% agar solution mixed with culture medium was used to form the bottom layer in 12-well plates. After solidification, 1.3 × 10^3^ cells were suspended in a mixture of agar solution and either control medium or S4-containing medium (at the 24-hour IC_50_ value) and added as the top layer. Culture medium (either with or without S4) was added, and cells were incubated for 30 days. Colonies were imaged using a bright-field microscope (SOPTOP ICX41), and ImageJ’s “ParticleSizer” plugin was used for colony number and size quantification.

### Wound healing assay

The wound healing assay was performed to measure cell migration upon S4 treatment. Cells (1 × 10^5^ per well) were seeded in 24-well plates with 1 mL of culture medium and incubated for 24 hours. The medium was then replaced with S4-containing medium (24-h IC_50_ value for each cell) or control medium. Then, a 200 μL pipette tip was used to create a cell-free wound area. Bright-field microscope images (SOPTOP ICX41) were captured at 0, 6, and 24 hours. Wound closure was quantified using the MRI_Wound_Healing_Tool plugin in ImageJ software.

### Apoptosis assay

To assess the apoptotic effects of S4, the Alexa Fluor® 488 Annexin V/Dead Cell Apoptosis Kit (cat. no. V13242, Thermo Fisher Scientific) was utilized. BC cells (1 × 10^5^ per well) were seeded in 12-well plates with 1 mL of culture medium and incubated at 37°C in 5% CO_2_ for 24 hours. The medium was then replaced with either S4-containing medium (24-h IC_50_ values for each cell) or control medium. Following 24-hour incubation, cells were harvested, washed with cold phosphate-buffered saline (PBS), and resuspended in 100 μL of 1X annexin-binding buffer. FITC Annexin V and propidium iodide (PI) working solution were added to each sample, followed by 15 minutes of incubation at room temperature. After adding 400 μL of 1X annexin-binding buffer, apoptotic cell populations were analyzed using flow cytometry (BD Accuri™ C6 Plus).

### Proliferation assay

Cell proliferation following S4 treatment was evaluated using the Click-iT™ EdU Cell Proliferation Kit (cat. no. C10337, Thermo Fisher Scientific). A total of 10,000 cells per well were plated in 96-well plates containing 100 μL of culture medium and incubated overnight at 37°C. After 24 hours of incubation with S4-containing medium or culture medium, 10 μL of EdU solution was added to each well, and cells were further incubated for 2 hours. Cells were then fixed with 100% methanol, washed with bovine serum albumin in PBS, and permeabilized using a saponin-based reagent. The Click-iT® reaction cocktail was prepared according to the manufacturer’s protocol and applied to the wells for EdU detection. After incubation at room temperature, cells were stained with Hoechst dye, and images were captured using a fluorescence microscope (Nikon Eclipse Ts2). The number of EdU-positive and Hoechst-positive cells was quantified using ImageJ software.

### Cell-cycle analysis

Propidium iodide (PI) staining was performed to analyze the cell cycle profile of control and S4-treated cells. BC cells (1 × 10^5^ per well) were plated in 12-well plates with 100 μL of culture medium and incubated for 24 hours. The medium was then replaced with either S4-containing or control medium, and incubation continued for an additional 24 hours. After treatment, cells were harvested, washed with PBS, and fixed in 70% ethanol for 30 minutes at 4°C. Following fixation, cells were washed twice with PBS and treated with 50 μL of a 100 μg/mL RNase A solution for 30 minutes at 37°C to prevent RNA interference in PI staining. Finally, 200 μL of PI solution (50 μg/mL) was added to each well, and the samples were analyzed via flow cytometry (BD Accuri™ C6 Plus) to determine cell cycle distribution.

### Gene expression analysis

To investigate the molecular mechanisms underlying the observed cellular effects of S4 treatment, quantitative real-time PCR (qRT-PCR) was performed on a panel of proto-oncogenes and tumor suppressor genes as previously listed before [14]. BC cells (3 × 10^5^ per well) were seeded in 6-well plates containing 2 mL of culture medium and incubated for 24 hours. Following incubation, the medium was replaced with either fresh culture medium or S4-containing medium, and cells were further incubated for 24 hours. Total RNA was extracted using a previously described protocol [14]. The isolated RNA was then reverse transcribed into cDNA, and qRT-PCR was carried out using gene-specific primers. GAPDH was used as the housekeeping gene for normalization, and relative gene expression was calculated using the 2^−^ΔΔCt method.

### Metabolome profiling

To examine the metabolic alterations induced by S4, untargeted metabolomics analysis was performed, as previously described [10]. Briefly, BC cells (3 × 10^5^ per well) were cultured in 6-well plates and incubated for 24 hours. The medium was then replaced with either control medium or S4-containing medium at the 24-hour IC_50_ concentration for each cell, and cells were incubated for an additional 24 hours. After treatment, cells were harvested by trypsinization, transferred to 5 mL centrifuge tubes, and centrifuged at 1,200 rpm for 5 minutes. The cell pellets were washed three times with prechilled PBS, followed by the addition of PBS in a 9:1 ratio. After another round of centrifugation, 200 μL of an extraction solution (containing d-Valine and d-Phenylalanine in Acetonitrile:Methanol:Acetone, 8:1:1 ratio) was added, and samples were vortexed thoroughly. Five cycles of sonication were performed to ensure cell lysis, with samples kept on ice between cycles. For protein precipitation, the samples were incubated at -20°C for 1 hour, followed by centrifugation at maximum speed for 15 minutes in a refrigerated centrifuge. The resulting supernatant was transferred to a clean plate, and the solvent was evaporated under nitrogen. Samples were then reconstituted in 50 μL of water, transferred to vials, and analyzed using an ultra-high-performance liquid chromatography/mass spectrometry (UHPLC-MS) system. The chromatographic conditions were as follows: column temperature: 45°C, sample chamber temperature: 8°C, and injection volume: 1 μL. Mobile phase A consisted of ultrapure water with 0.1% formic acid, while mobile phase B contained methanol with 0.1% formic acid. A 30-minute gradient was applied at a flow rate of 0.3 mL/min. Electrospray ionization (ESI) was performed in both positive (ESI^+^) and negative (ESI^−^) ionization modes in separate runs. MSE acquisition mode was used to collect both primary and secondary mass spectrometry data. Peak detection, calibration, and normalization of raw LC-MS data were conducted using Compound Discoverer 3.3 SP1 (Thermo Scientific) software. Metabolite identification was performed using ChemSpider, Human Metabolome Database, mzCloud, and mzVault databases. The fold change (FC) threshold for significantly altered metabolites was set at >1.5, with a p-value <0.05. Pathway enrichment analysis and metabolic pathway processing were performed using MetaboAnalyst 6.0 [15] and the KEGG compound database. Differential metabolites were visualized using the EnhancedVolcano package in R.

### Statistical analysis

All experimental data are presented as mean ± standard deviation. Normality of data distribution was assessed using the Shapiro–Wilk test. The One-Way ANOVA test, followed by Tukey’s post hoc test, was used for statistical comparisons in the cell viability assay. For all other assays, significance was determined using a Student’s t-test. A p-value < 0.05 was considered statistically significant. Statistical analyses and data visualization were performed using the ‘Statix’ package in R.

## Results

### S4 limits the growth of BC cells

To asses the impact of S4 on BC cells, we investigated their viability following treatment with various doses of S4 for 24 and 48 hours. A significant reduction in cellular viability was observed when the S4 concentration exceeded 0.01 mM in MCF-7 cells at both time points. At 0.4 mM S4, cells were completely non-viable (Figure 1A, B). Treatment with 0.05 mM S4 reduced MCF-7 viability to 73% and 61% at 24 and 48 hours, respectively (p-value < 0.001). At 0.1 mM S4, viability decreased further to 47% and 32% at the respective time points (p-value < 0.001). Exposure to 0.2 mM S4 resulted in viability reductions to 22% and 11% after 24 and 48 hours, respectively (p-value < 0.001). The IC50 values for MCF-7 cells were calculated as 0.094 mM at 24 hours and 0.067 mM at 48 hours.

**Figure 1.**
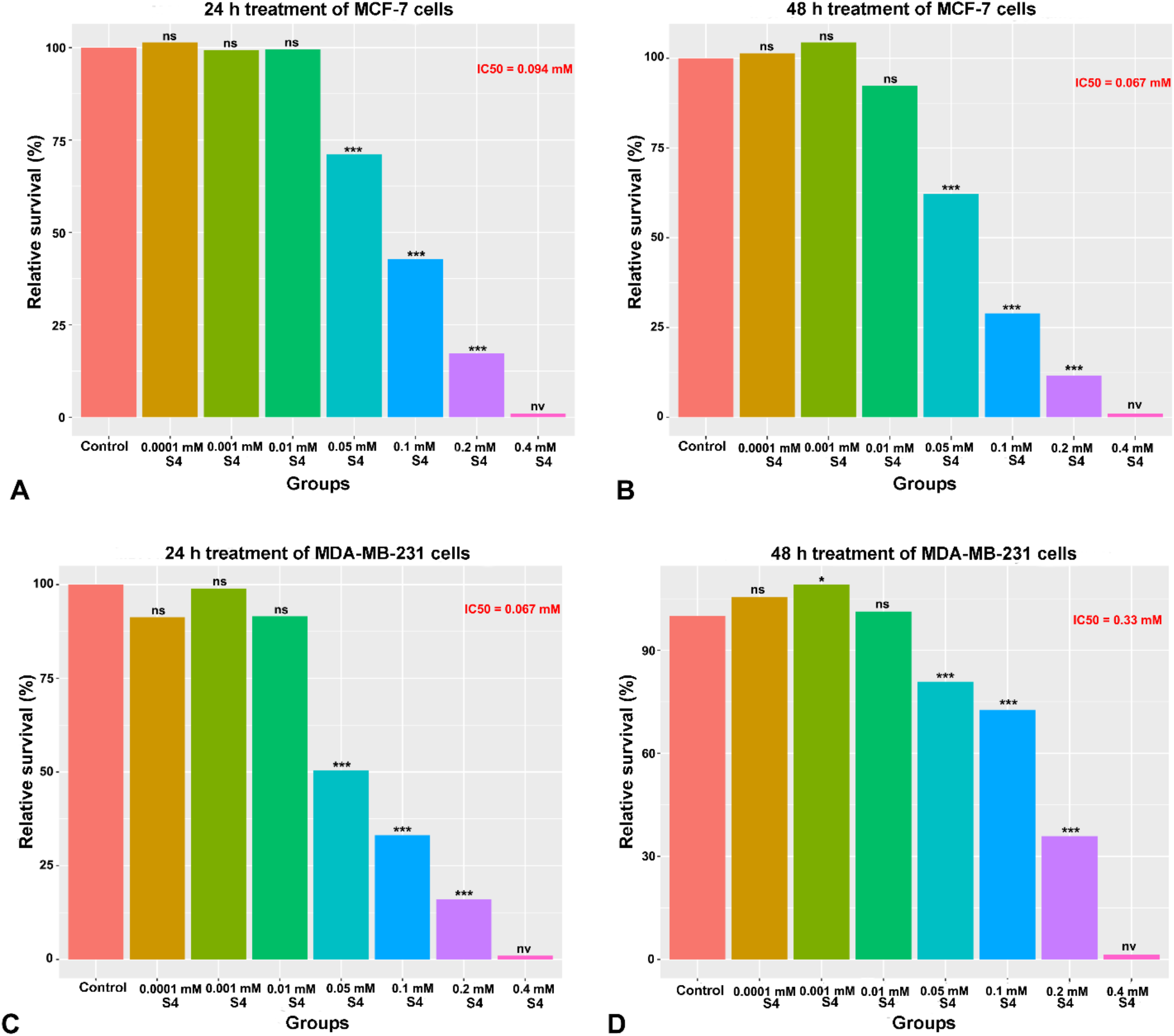
S4 suppresses BC viability in a dose dependent manner. Relative viability of MCF-7 cells upon S4 administration along **A)** 24 and **B)** 48 hours. **C)** 24 hours and **D)** 48 hours S4-treated MDA-MB-231 viability. ***p-value < 0.001, **p-value < 0.01, *p-value < 0.05. ns; non-significant. nv; non-viable.

A similar trend was observed in MDA-MB-231 cells (Figure 1C, D). Treatment with 0.05 mM S4 diminished cell viability to 83% and 51% at 24 and 48 hours, respectively (p-value < 0.001). Exposure to 0.1 mM S4 resulted in 78% and 32% viability, while 0.2 mM S4 decreased viability to 36% and 17% at the respective time points (p-value < 0.001). The IC50 values for MDA-MB-231 cells were determined as 0.067 mM for 24 hours and 0.033 mM for 48 hours.

The observed effects of S4 on cell viability prompted us to investigate its impact on BC cell clonogenicity. In MCF-7 cells, all tested S4 concentrations (0.025 mM, 0.05 mM, and 0.1 mM) significantly inhibited colony formation, with a 1.5-fold reduction at 0.025 mM (p-value < 0.01), a 2.1-fold reduction at 0.05 mM (p < 0.01), and a 3.6-fold reduction at 0.1 mM (p-value < 0.001) (Figure 2A). Similarly, treatment with 0.1 mM S4 markedly impaired the clonogenic potential of MDA-MB-231 cells, reducing colony formation by 1.6-fold compared to control wells (p-value < 0.001) (Figure 2B).

**Figure 2.**
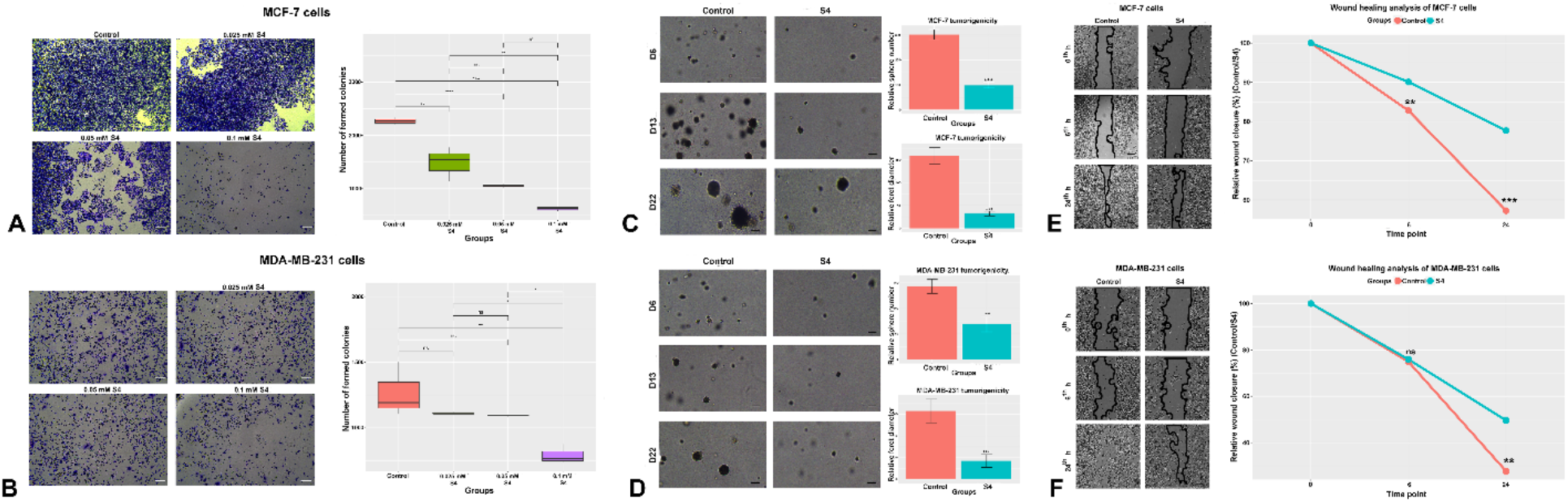
S4 impairs clonogenicity of BC cells and inhibits their migration. **A)** The representative images and boxplot graphs of MCF-7 colonies following S4 treatment. **B)** Representative images of S4-treated and control MDA-MB-231 colonies and colony formation analysis of MDA-MB-231 cells. Relative spheres of **C)** MCF-7 and **D)** MDA-MB-231 cells after S4 treatment. The representative 24-hour migration images and wound healing analysis of **E)** MCF-7 and **F)** MDA-MB-231 cells. ***p-value < 0.001, **p-value < 0.01, *p-value < 0.05. ns; non-significant D; day. H; hour. Scale bar; 50 μm.

To determine the effect of S4 on tumorigenicity, we evaluated the anchorage-independent spheroid formation capacity of BC cells (Figure 2C, D). S4 treatment significantly decreased both the mean spheroid number and feret diameter at the examined time points in both cell lines. In MCF-7 cells, spheroid numbers and feret diameters were reduced by 3-fold and 3.6-fold, respectively (p-value < 0.001). In MDA-MB-231 cells, a 2.4-fold reduction in spheroid number (p-value < 0.01) and a 3-fold reduction in feret diameter (p-value < 0.001) were observed. These findings demonstrate that S4 inhibits BC cell clonogenicity through both anchorage-dependent and anchorage-independent mechanisms, further supporting its growth-limiting effects on BC cells.

### S4 substantially inhibits BC cell migration

The migratory capacity of BC cells was examined using a wound healing assay, revealing a significant anti-migratory effect of S4. In MCF-7 cells, wound closure was reduced by 1.2-fold at the 6-h time point and by 2-fold at 24 hours (p-value < 0.01 for 6 h, p-value < 0.001 for 24 h) (Figure 2E). Likewise, in MDA-MB-231 cells, wound closure decreased by 2.6-fold at 24 hours (p < 0.01), indicating a pronounced inhibition of cell migration (Figure 2F).

### S4 induces apoptosis while inhibiting proliferation and arresting the cell-cycle arrest in BC cells

Given the substantial cytotoxicity and growth-inhibitory effects of S4, we sought to uncover the molecular mechanisms underlying these phenomena. To this end, we analyzed apoptosis, proliferation, and cell cycle progression following S4 treatment (Figure 3). Our findings demonstrated that S4 significantly increased apoptotic activity in MCF-7 and MDA-MB-231 cells, with a 1.6- and 2-fold rise in apoptotic cell numbers, respectively (p-value < 0.05) (Figure 3A, B).

**Figure 3.**
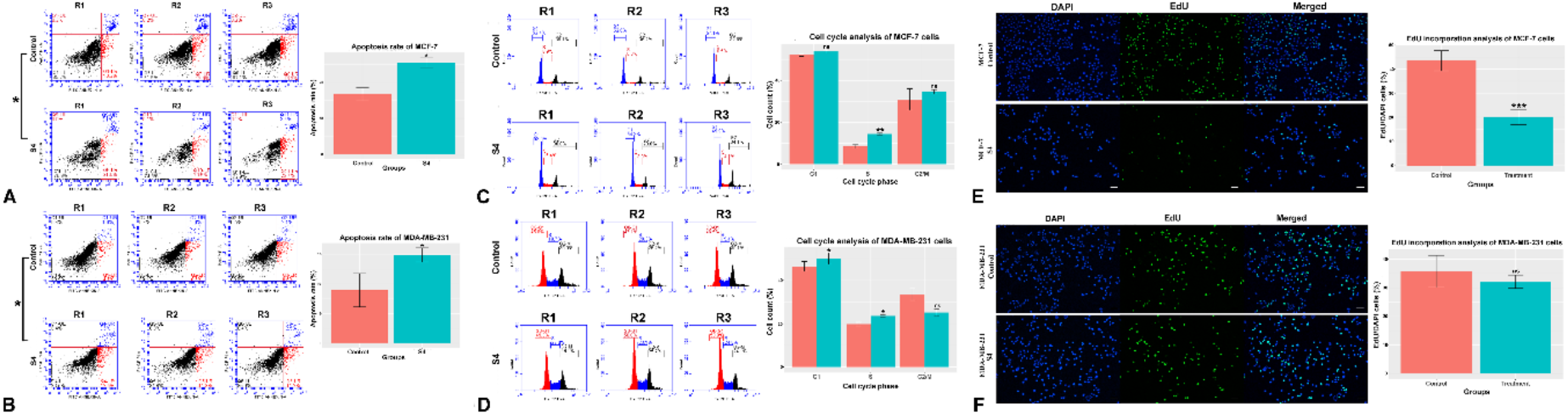
S4 induces apoptosis and cell cycle arrest in BC cells while inhibiting MCF-7 proliferation. The **A)** MCF-7 and **B)** MDA-MB-231 diagrams following S4 treatment. The right upper quadrant represents (the blue dots) late apoptotic cells. The right lower quadrant demonstrates (the red dots) early apoptotic cells. The left upper quadrant shows necrotic cells, and the left lower quadrant shows live cells. The flowcytometry charts of **C)** MCF-7 and **D)** MDA-MB-231 cells. The EdU incorporation of **E)** MCF-7 and **F)** MDA-MB-231 cells. *p-value < 0.05. ***p-value < 0.001. ns; non-significant. PerCP; peridinin-chlorophyll-protein channel. FITC; Fluorescein isothiocyanate channel. R; replicate. DAPI; 4’,6-diamidino-2-phenylindole. EdU; 5–ethynyl–2′– deoxyuridine. Scale bar; 50 μm.

Beyond its pro-apoptotic effects, S4 also exerted a profound impact on cell cycle regulation. In MCF-7 cells, S4 induced significant S-phase arrest, leading to a 2.4-fold decrease in cell cycle progression (p-value < 0.01) (Figure 3C). In MDA-MB-231 cells, S4 caused cell cycle arrest at both the G0/G1 and S phases, resulting in a 1.5-fold reduction in cell cycle progression (p-value < 0.05) (Figure 3D). Additionally, the EdU incorporation assay confirmed a significant reduction in proliferation, with a 2.1-fold decrease in the proportion of EdU-positive cells in MCF-7 cells after S4 treatment (p-value < 0.001) (Figure 3E). No significant changes in proliferation were detected in MDA-MB-231 cells (Figure 3F). Collectively, these findings suggest that S4 promotes apoptosis and induces cell cycle arrest in both BC cell lines while selectively inhibiting proliferation in MCF-7 cells.

### S4 modulates gene expression in an anti-carcinogenic manner

To further elucidate the molecular changes induced by S4, we performed gene expression analysis in S4-treated cells (Table 1). In MCF-7 cells, S4 treatment significantly upregulated *GADD45A* (3.45-fold, p-value < 0.01), CDKN1B (2.42-fold, p-value < 0.001), and *PUMA* (3.28-fold, p-value < 0.01), while downregulating *ANKRD1* (1.5-fold decrease, p-value < 0.01), *CXCL1* (18-fold decrease, p-value < 0.05), *EDN1* (1.5-fold decrease, p-value < 0.05), *CDK6* (1.59-fold decrease, p-value < 0.01), *CCND1* (1.5-fold decrease, p-value < 0.05), *ATM* (2.19-fold decrease, p-value < 0.05), and *MMP9* (7.6-fold decrease, p-value < 0.01) (Table 1)

**Table 1.**
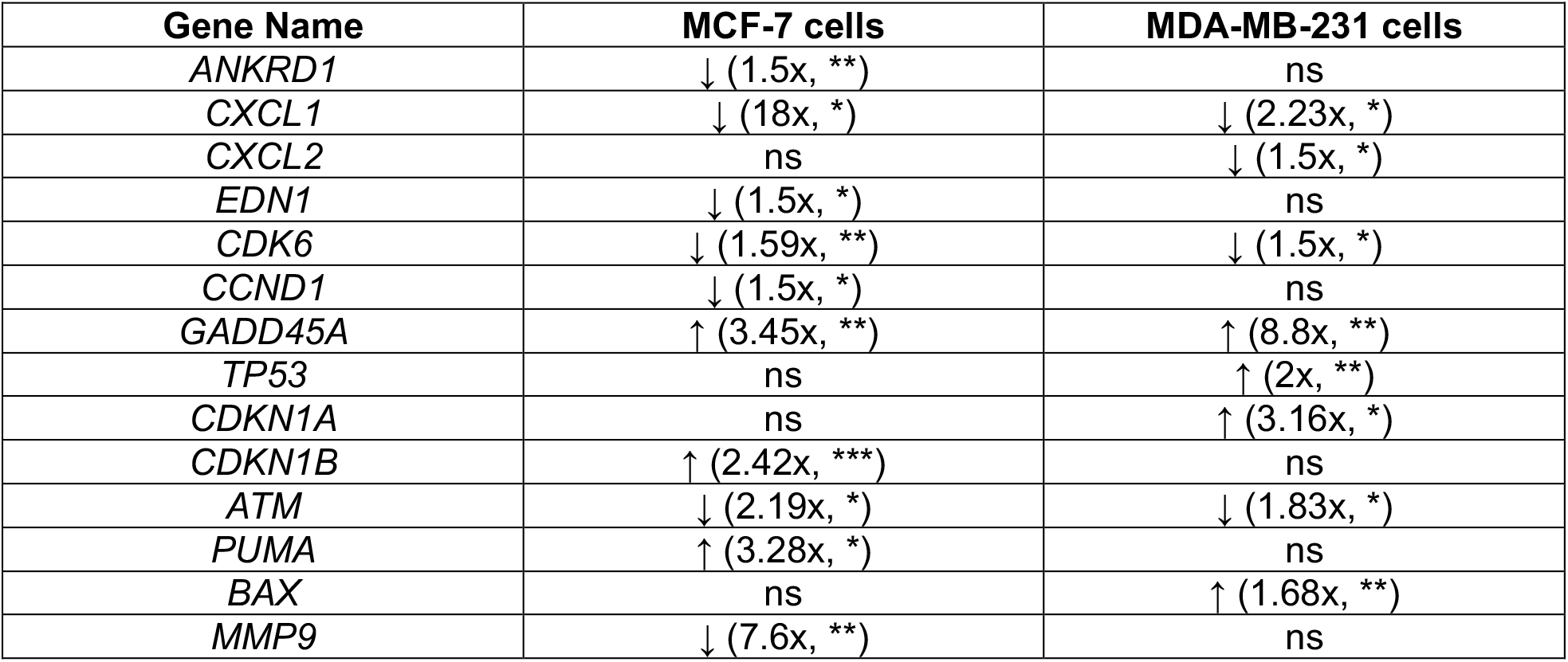
Gene expression profile of BC cells. ↓; downregulated. ↑; upregulated. ***p-value < 0.001. **p-value < 0.01. *p-value < 0.5. ns; non-significant.

Similarly, in MDA-MB-231 cells, S4 treatment led to significant upregulation of *GADD45A* (8.8-fold, p-value < 0.01), *TP53* (2-fold, p-value < 0.01), *CDKN1A* (3.16-fold, p-value < 0.05), and *BAX* (1.68-fold, p-value < 0.01), along with downregulation of *CXCL1* (2.23-fold decrease, p-value < 0.05), *CXCL2* (1.5-fold decrease, p-value < 0.05), *CDK6* (1.5-fold decrease, p-value < 0.05), *CCND1* (1.5-fold decrease, p-value < 0.05), and *ATM* (1.83-fold decrease, p-value < 0.05) (Table 1).

Taken together, these results indicate that S4 activates genes associated with apoptosis and cell cycle arrest while repressing genes involved in proliferation and survival, reinforcing its anti-carcinogenic potential.

### S4 stimulates significant metabolic changes in BC cells

Significant metabolic differences between control and treatment groups are consistently demonstrated by the metabolome profile and differential metabolite analysis. Differential expression of metabolites in S4-treated BC cells compared to control conditions is displayed in a volcano plot (Figure 4A, B). Major metabolites up-regulated in MCF-7 cells include 6-oxohexanoic acid, galidesivir, D-(-)-glutamine, 2-mercaptoethanol, and 4-methyl-2-oxo-2H-chromene-7-YL 5-(acetylamino)-3,5-dideoxy-L-erythro-NON-2-ulopyranosidonic acid (Figure 4A); and paracetamol, 4-pyridoxic acid, 6-quinolonecarboxylic acid, 5-methylcytidine and 8-fluoro-4H-thiochromono[3,4-d]isoxazole, highlighting significant metabolic alteration upon S4 treatment in MDA-MB-231 cells (Figure 4B).

**Figure 4.**
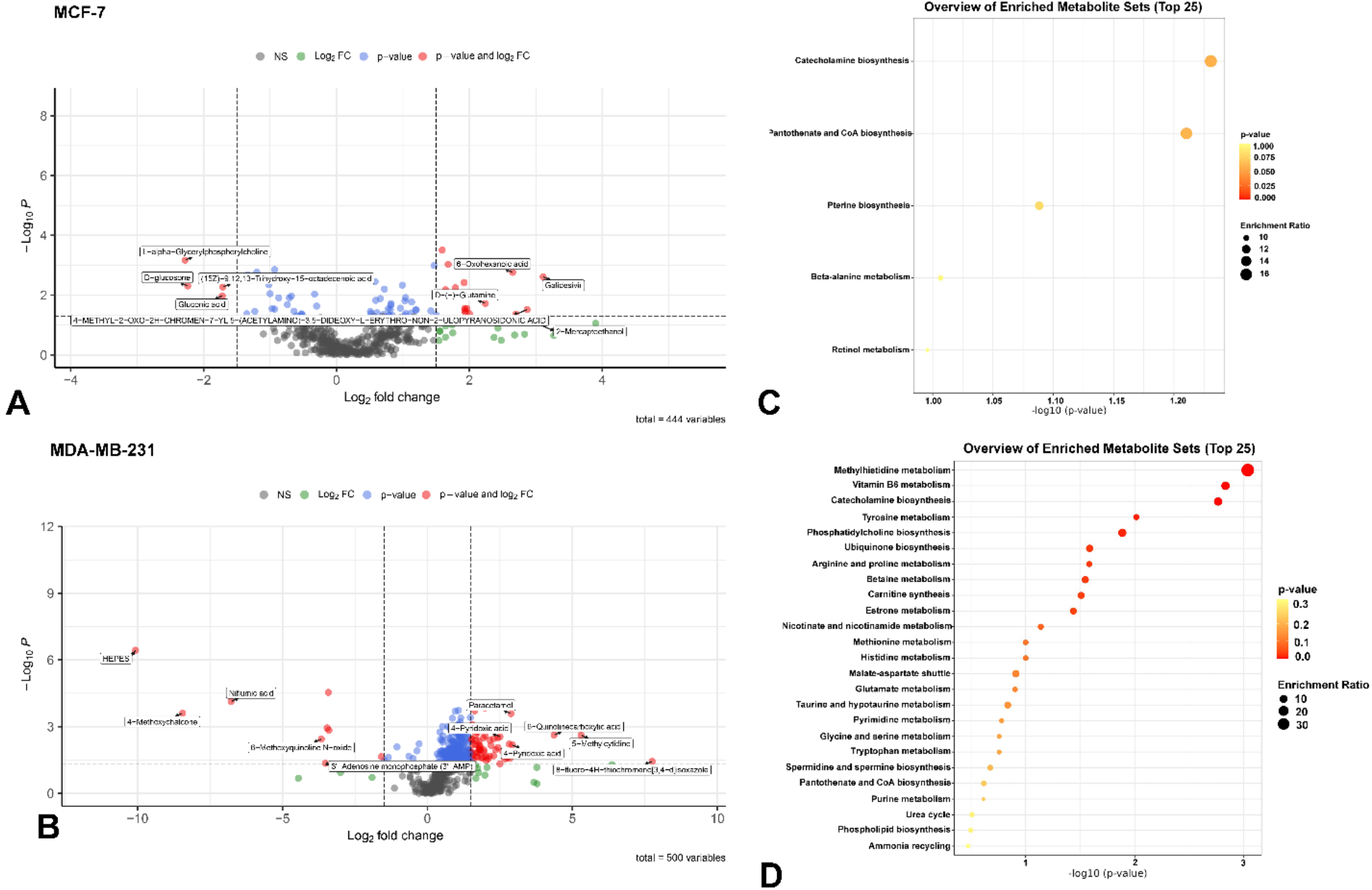
S4 induces metabolic reprogramming in BC cells. Identified top five upregulated and downregulated metabolites in **A)** MCF-7 and **B)** MDA-MB-231 cells. Gray dots represent non-significant metabolites. Green dots represent the statistically significant up- or down-regulated metabolites in terms of log2 fold-change. Blue dots represent statistically significant up- or down-regulated metabolites in terms of p-value. Red dots represent statistically significant up- or down-regulated metabolites in terms of both log2 fold-change and p-value. ns; non-significant. FC; fold-change. **C)** MCF-7 and **D)** MDA-MB-231 enrichment plots. The dot plot displays the top 25 enriched metabolite sets identified through pathway analysis. The size of each dot reflects the enrichment ratio, with larger dots indicating higher values, while the color represents the corresponding p-value.

Besides, L-alpha-Glycerylphosphorylcholine, D-glucoson, (15Z)-9,12,13-Trihydroxy-15-octadecenoic acid, Tetrahydro-2,5-furan-diacetic acid and Gluconic acid in MCF-7 cells (Figure 4A) and HEPES, Niflumic acid, 4-Methoxychalcone, 6-Methoxyquinoline N-oxide and 3’-Adenosine monophosphate (3’-AMP) in MDA-MB-231 cells were identified as the most significantly down-regulated metabolites following S4 administration (Figure 4B).

Metabolite set enrichment analysis provided deeper insights into the mechanism of action of S4. Both cell lines exhibited alterations in the catecholamine biosynthetic pathway. In S4-treated MDA-MB-231 cells, significant metabolic changes were observed in amino acid, nucleotide, and energy metabolism, including tyrosine metabolism, arginine metabolism, ubiquinone metabolism, and methionine metabolism (Figure 4D). Additionally, tryptophan metabolism and ammonia recycling were among the prominently enriched pathways in these cells (Figure 4C, D).

## Discussion

BC is the second most common malignancy worldwide, primarily affecting women [16,17]. Its pathogenesis involves genetic and environmental factors, with aging and hormonal changes increasing the risk of relapse and mortality. BC is classified by hormone receptor expression (Estrohen receptor -ER-, progesterone receptor -PR-, Human epidermal growth factor receptor 2 -HER2-), guiding treatment strategies such as tamoxifen and aromatase inhibitors for hormone receptor-positive cases [18]. However, triple negative breast cancer (TNBC), lacking these receptors, requires alternative therapies, with AR emerging as a promising target [19–22].

It has been demonstrated that modulating AR signaling can suppress BC growth both *in vitro* and *in vivo* [23,24]. Given the severe side effects associated with androgen-based therapies [25,26], SARMs have gained attention as potential alternatives [8,27]. Although the anti-tumorigenic impact of enzalutamide, a nonsteroid antiandrogen used in prostate cancer treatment [28], and Enobosarm (Gtx-024), a non-steroidal SARM [29], have been investigated clinically for ER+ BC and TNBC patients [30–32], the impact of S4, a member of relatively new class of SARM [33], has not been previously investigated in BC before. This study provides the first evidence of S4’s anti-neoplastic effects in AR+ ER+ and TNBC cell lines.

Our findings demonstrate that S4 exerts significant anti-carcinogenic effects on BC cells by inhibiting proliferation, migration, and clonogenic potential while promoting apoptosis and cell cycle arrest. The dose-dependent reduction in cell viability observed in both MCF-7 and MDA-MB-231 cells suggests a potent cytotoxic effect of S4, with IC50 values indicating higher sensitivity of MDA-MB-231 cells compared to MCF-7 cells. This differential sensitivity may stem from intrinsic molecular variations between these cell lines, such as differences in hormone receptor status and genetic background. The strong anti-carcinogenic potential of S4 in various cancer types has been documented in earlier reports [10–13]. In non-small cell lung cancer, S4 not only inhibited proliferation, migration, and survival but also significantly altered the metabolome profile of A549 cells [10]. Metabolomic analysis revealed notable shifts in key metabolic pathways, particularly those involved in energy metabolism, amino acid biosynthesis, and oxidative stress regulation. These metabolic changes enriched cancer-associated pathways, indicating that S4 influences A549 cells by restructuring their metabolic landscape [10]. In hepatocellular carcinoma, S4 suppressed the PI3K/AKT/mTOR pathway—a key driver of tumor progression and poor prognosis [11]. In pancreatic cancer, S4 exerted a cytostatic influence by inducing G0/G1 cell cycle arrest, effectively halting proliferation [12]. In both temozolomide-sensitive and -resistant glioblastoma multiforme cells, S4 suppressed cell growth and migration while promoting apoptosis, reactive oxygen species generation, and cellular senescence [13]. Gene expression analysis further revealed that S4 exposure upregulated apoptosis-, DNA damage-, and senescence-associated genes while downregulating those linked to cellular proliferation [13].

Previous studies demonstrated that SARMs, including Gtx-024 and Gtx-027, suppressed TNBC viability *in vitro* and tumor growth *in vivo* [34]. In a patient-derived AR+/ER+ BC xenograft model, RAD-140, another SARM, specifically bound to AR in breast tissue and inhibited tumor growth while blocking estrogen signaling via *ESR1* [35]. Furthermore, in ex-vivo patient-derived explant model, DHT suppressed the cell proliferation by neutralizing the tumor-promoting effect of estrogen [36] which aligns well with our EdU-staining data where S4 inhibited the proliferation of BC cells. Similarly, SARMs such as Gtx-024, Gtx-027, SARM-2F, and T8039 have been shown to modulate AR signaling and inhibit prostate cancer growth both *in vitro* and *in vivo* [37]. The clonogenic assay results further reinforce the ability of S4 to suppress long-term cell survival, as evidenced by the dose-dependent reduction in colony formation in both cell lines. The impairment of anchorage-independent growth observed in spheroid formation assays highlights S4’s potential in inhibiting tumorigenicity. Given that anchorage-independent growth is a hallmark of malignant transformation, these results suggest that S4 may disrupt key pathways essential for tumor progression.

S4 treatment induced apoptosis while arresting the cell cycle at different phases in the two cell lines. The observed S-phase arrest in MCF-7 cells and combined G0/G1-S phase arrest in MDA-MB-231 cells align with the downregulation of key cell cycle regulators, including *CDK6* and *CCND1*. These findings suggest that S4 disrupts the normal progression of the cell cycle, leading to decreased proliferation. The selective inhibition of proliferation in MCF-7 cells, as evidenced by the EdU incorporation assay, may be linked to their hormone receptor-positive status, potentially indicating a unique dependency on proliferative signaling pathways that S4 may target. Interestingly, while MDA-MB-231 cells did not show a significant reduction in proliferation *via* the EdU assay, the observed G0/G1 arrest suggests that these cells were unable to transition into the S phase. This could explain the lack of EdU incorporation, as the assay specifically labels cells undergoing DNA synthesis. Therefore, additional assays that directly mark cell division, such as Ki-67 staining or phospho-histone H3 detection, could provide further clarification on the proliferative status of MDA-MB-231 cells following S4 treatment.

Inhibition of cell migration is another crucial finding, as metastasis remains a major challenge in BC treatment. The wound healing assay showed a significant decrease in cell motility following S4 treatment, with MDA-MB-231 cells displaying a more notable reduction. This is particularly significant given the highly invasive nature of MDA-MB-231 cells, suggesting that S4 may have a stronger effect on aggressive BC phenotypes. This migration-inhibitory effect of S4 is consistent with previous reports demonstrating similar outcomes in hepatocellular carcinoma cells [11]. Further investigations into molecular mechanisms underlying this anti-migratory effect—such as changes in epithelial-mesenchymal transition (EMT) markers—could provide additional insights.

To further explore the molecular mechanisms underlying S4’s anti-neoplastic effects, we conducted qRT-PCR analyses to identify its molecular targets in BC cells. In MCF-7 cells, S4 treatment led to the significant downregulation of *ANKRD1*, a mesenchymal-specific transcriptional coregulator known to drive cancer-associated fibroblast (CAF) conversion [38]. The silencing of *ANKRD1* in CAFs has been shown to suppress sphere formation and invasion in SCC13 and FaDu cells, as well as tumor growth *in vivo* [38]. Moreover, *CXCL1*, a key driver of migration and epithelial-to-mesenchymal transition in BC via MAPK activation [39], was downregulated following S4 treatment, supporting its observed anti-migratory effects. Additionally, *EDN1*, which promotes BC proliferation and migration via the AKT pathway [40,41] was significantly downregulated in S4-treated MCF-7 cells, further corroborating its anti-proliferative and anti-migratory properties. The suppression of *CDK6, CCND1*, and *MMP-9* upon S4 treatment reinforces these findings, while the upregulation of *GADD45A, PUMA*, and *CDKN1B* suggests enhanced apoptosis and cell cycle arrest [42].

In MDA-MB-231 cells, S4 treatment similarly resulted in the downregulation of oncogenic factors such as *CDK6, ATM, CXCL1*, and *CXCL2*. Given that *CDK6* plays a crucial role in cell cycle progression and oncogenesis [43], its suppression provides additional support for S4’s anti-neoplastic activity. *ATM*, a key regulator of DNA damage response, has been implicated in BC progression and cancer stem cell maintenance [44–46]. Its downregulation following S4 treatment suggests a novel therapeutic avenue for targeting ATM in BC. Furthermore, the upregulation of *GADD45A, TP53, CDKN1A*, and *BAX*, which regulate apoptosis and cell cycle arrest [47–50] provides further mechanistic insight into S4’s anti-tumorigenic effects.

Changes in metabolism have lately been identified as one of the characteristics of cancer. Cancer cells have high biosynthetic and bioenergetic demands since they rapidly replicate and proliferate [51]. In MCF-7 cells, S4 treatment downregulated L-alpha-Glycerylphosphocholine, a key metabolite in choline metabolism, which is often elevated in cancer and considered an oncometabolic factor [52], and the downregulation of above-mentioned metabolite further layers another evidence into S4’s anti-neoplastic activity in BC. Additionally, S4 reduced D-glucosone, suggesting potential interference with glucose metabolism, a hallmark of cancer progression [53– 55].

In MDA-MB-231 cells, S4 increased 6-quinolinecarboxylic acid, a compound with known anti-cancer properties [56,57]. It also downregulated 3-adenosine monophosphate (3-AMP), which may affect cyclic AMP signaling, reducing tumorigenic factors such as cyclin-D1, β-catenin, and c-myc while potentially altering ERK1/2, JNK, and p38 pathways [58]. Downregulation of 3-AMP upon S4 administration might be related to its receptor and need to be deciphered in future studies. Substantial metabolic alterations following S4 treatment were also evident in *in-silico* pathway analysis. S4 treatment significantly altered key metabolic pathways, including amino acid, nucleotide, and energy metabolism, disrupting essential processes required for cancer cell survival and proliferation. In MDA-MB-231 cells, it affected phosphatidylcholine biosynthesis, glucose metabolism, and oxidative stress-related pathways, potentially impacting membrane lipid composition and energy production. The downregulation of key metabolites in these pathways aligns with the observed reduction in cell viability and proliferation. Similarly, in MCF-7 cells, S4 influenced catecholamine biosynthesis, tryptophan metabolism, and ammonia recycling, suggesting an effect on stress response mechanisms and metabolic adaptation. Alterations in nucleotide metabolism and redox homeostasis further support the hypothesis that S4 disrupts critical metabolic networks essential for tumor growth and metastasis. These findings suggest that S4 exerts anti-tumorigenic effects by reprogramming metabolic pathways essential for BC cell survival and progression, thereby inhibiting proliferation, migration, and tumorigenicity.

This study has potential limitations. While S4 demonstrates efficacy in AR+/ER+ and TNBC cell lines, the precise role of AR signaling remains unclear. Future studies should distinguish AR-dependent and AR-independent mechanisms through genetic and pharmacological approaches, such as AR knockdown or competitive inhibition. To validate S4’s therapeutic potential, in vivo studies using xenograft and patient-derived tumor models are necessary. Moreover, evaluating its toxicity and off-target effects in animal models will be crucial for clinical translation. Regarding S4’s impact on multiple oncogenic pathways, combination therapies with existing BC treatments— including endocrine therapy (tamoxifen, aromatase inhibitors), chemotherapy (cisplatin, doxorubicin), and targeted therapy (CDK4/6 and PARP inhibitors)—should be explored. Investigating potential synergistic or additive effects could provide an enhanced therapeutic strategy, particularly for drug-resistant BC subtypes.

## Conclusion

This study provides the first evidence that S4, a previously underexplored SARM, exerts significant anti-carcinogenic effects in BC cells. S4 effectively reduces cell viability, suppresses clonogenicity and migration, induces apoptosis, and causes cell cycle arrest in both MCF-7 and MDA-MB-231 cells. Gene expression analysis further supports these effects, revealing the upregulation of tumor suppressor genes and reinforcing S4’s role in inhibiting tumor progression. Untargeted metabolomics analysis highlights S4-induced metabolic reprogramming, with significant alterations in phosphatidylcholine biosynthesis, catecholamine biosynthesis, and nucleotide metabolism—pathways crucial for cancer cell survival. These findings suggest that S4’s anti-tumor activity extends beyond AR signaling modulation, positioning it as a promising candidate for BC therapy. Given these promising *in vitro* effects, future studies should validate S4’s efficacy in in vivo models, assess its potential synergy with existing BC treatments, and evaluate its long-term safety profile. This study lays the groundwork for further exploration of SARMs in BC treatment, offering a novel avenue for targeted therapy.

## Author Contributions

Turan Demircan concepted and designed the study. Material preparation and data collection were perfomed by Mervenur Yavuz and Turan Demircan. Data analyses were performed by Mervenur Yavuz and Turan Demircan. The first draft of the manuscript was written by Mervenur Yavuz and edited by Turan Demircan. All authors read and approved the final manuscript.

## Ethics Approval and Consent to Participate

Not applicable.

## Consent for Publication

Not applicable.

## Availability of Data and Materials

The data and supportive information are available within the article.

## Acknowledgements

Declared none.

## References

1. Lukasiewicz S, Czeczelewski M, Forma A, Baj J, Sitarz R, Stanisławek A. Breast Cancer— Epidemiology Risk Factors, Classification, Prognostic Markers, and Current Treatment Strategies—An Updated Review. Cancers (Basel). 2021 Aug 25;13(17):4287.

2. Swain SM, Shastry M, Hamilton E. Targeting HER2-positive breast cancer: advances and future directions. Nature Reviews Drug Discovery. 2022 Nov 7;22(2):101.

3. Dart DA, Bevan CL, Uysal-Onganer P, Jiang WG. Analysis of androgen receptor expression and activity in the mouse brain. Sci Rep. 2024 May 15;14(1):11115.

4. Wen S, Niu Y, Huang H. Posttranslational regulation of androgen dependent and independent androgen receptor activities in prostate cancer. Asian Journal of Urology. 2020 Jul 1;7(3):203–18.

5. Dehm SM, Tindall DJ. Ligand-independent Androgen Receptor Activity Is Activation Function-2-independent and Resistant to Antiandrogens in Androgen Refractory Prostate Cancer Cells*. Journal of Biological Chemistry. 2006 Sep 22;281(38):27882–93.

6. Anestis A, Zoi I, Papavassiliou AG, Karamouzis MV. Androgen Receptor in Breast Cancer— Clinical and Preclinical Research Insights. Molecules. 2020 Jan;25(2):358.

7. Narayanan R, Coss CC, Dalton JT. Development of selective androgen receptor modulators (SARMs). Molecular and Cellular Endocrinology. 2018 Apr 15;465:134–42.

8. Christiansen AR, Lipshultz LI, Hotaling JM, Pastuszak AW. Selective androgen receptor modulators: the future of androgen therapy? Transl Androl Urol. 2020 Mar;9(Suppl 2):S135–48.

9. Kearbey JD, Gao W, Fisher SJ, Wu D, Miller DD, Dalton JT. Ekects of Selective Androgen Receptor Modulator (SARM) Treatment in Osteopenic Female Rats. Pharm Res. 2009 Nov 1;26(11):2471– 7.

10. Demircan T, Yavuz M, Bölük A. Unveiling the Potential of S4 on Non-small Cell Lung Cancer Cells: Impact on Proliferation, Apoptosis, Senescence, and Metabolome Profile. http://www.eurekaselect.com [Internet]. [cited 2025 Mar 17]; Available from: https://www.eurekaselect.com/article/146020

11. Yavuz M, Takanlou LS, Avcı ÇB, Demircan T. A selective androgen receptor modulator, S4, displays robust anti-cancer activity on hepatocellular cancer cells by negatively regulating PI3K/AKT/mTOR signalling pathway. Gene. 2023 Mar 27;869:147390.

12. Bölük A, Yavuz M, Takanlou MS, Avcı ÇB, Demircan T. In vitro anti-carcinogenic ekect of andarine as a selective androgen receptor modulator on MIA-PaCa-2 cells by decreased proliferation and cell-cycle arrest at G0/G1 phase. Biochem Biophys Res Commun. 2023 Sep 3;671:132–9.

13. Yavuz M, Demircan T. Exploring the potentials of S4, a selective androgen receptor modulator, in glioblastoma multiforme therapy. Toxicology and Applied Pharmacology. 2024 Sep 1;490:117029.

14. Yavuz M, Demircan T. A potent ion channel blocker, hydroquinidine, exhibits strong anti-cancer activity on colon, pancreatic, and hepatocellular cancer cells. Mol Biol Rep [Internet]. 2023 Jan 12 [cited 2023 Feb 7]; Available from: https://doi.org/10.1007/s11033-023-08245-3

15. Pang Z, Lu Y, Zhou G, Hui F, Xu L, Viau C, et al. MetaboAnalyst 6.0: towards a unified platform for metabolomics data processing, analysis and interpretation. Nucleic Acids Research. 2024 Apr 8;gkae253.

16. Barzaman K, Karami J, Zarei Z, Hosseinzadeh A, Kazemi MH, Moradi-Kalbolandi S, et al. Breast cancer: Biology, biomarkers, and treatments. International Immunopharmacology. 2020 Jul 1;84:106535.

17. Kolak A, KamiŁska M, Sygit K, Budny A, Surdyka D, Kukiełka-Budny B, et al. Primary and secondary prevention of breast cancer. Ann Agric Environ Med. 2017 Dec 23;24(4):549–53.

18. Miller E, Lee HJ, Lulla A, Hernandez L, Gokare P, Lim B. Current treatment of early breast cancer: adjuvant and neoadjuvant therapy. F1000Res. 2014;3:198.

19. Dent R, Trudeau M, Pritchard KI, Hanna WM, Kahn HK, Sawka CA, et al. Triple-negative breast cancer: clinical features and patterns of recurrence. Clin Cancer Res. 2007 Aug 1;13(15 Pt 1):4429–34.

20. Doane AS, Danso M, Lal P, Donaton M, Zhang L, Hudis C, et al. An estrogen receptor-negative breast cancer subset characterized by a hormonally regulated transcriptional program and response to androgen. Oncogene. 2006 Jun 29;25(28):3994–4008.

21. McGhan LJ, McCullough AE, Protheroe CA, Dueck AC, Lee JJ, Nunez-Nateras R, et al. Androgen receptor-positive triple negative breast cancer: a unique breast cancer subtype. Ann Surg Oncol. 2014 Feb;21(2):361–7.

22. Qu Q, Mao Y, Fei X chun, Shen K wei. The Impact of Androgen Receptor Expression on Breast Cancer Survival: A Retrospective Study and Meta-Analysis. PLoS One. 2013 Dec 4;8(12):e82650.

23. Cochrane DR, Bernales S, Jacobsen BM, Cittelly DM, Howe EN, D’Amato NC, et al. Role of the androgen receptor in breast cancer and preclinical analysis of enzalutamide. Breast Cancer Res. 2014 Jan 22;16(1):R7.

24. Barton VN, D’Amato NC, Gordon MA, Lind HT, Spoelstra NS, Babbs BL, et al. Multiple molecular subtypes of triple-negative breast cancer critically rely on androgen receptor and respond to enzalutamide in vivo. Mol Cancer Ther. 2015 Mar;14(3):769–78.

25. Davey RA, Grossmann M. Androgen Receptor Structure, Function and Biology: From Bench to Bedside. Clin Biochem Rev. 2016 Feb;37(1):3–15.

26. Hartgens F, Kuipers H. Ekects of Androgenic-Anabolic Steroids in Athletes. Sports Med. 2004 Jul 1;34(8):513–54.

27. Gao W, Dalton JT. Expanding the therapeutic use of androgens via selective androgen receptor modulators (SARMs). Drug Discov Today. 2007 Mar;12(5–6):241–8.

28. Linder S, van der Poel HG, Bergman AM, Zwart W, Prekovic S. Enzalutamide therapy for advanced prostate cancer: ekicacy, resistance and beyond. Endocr Relat Cancer. 2018 Sep 14;26(1):R31–52.

29. Dalton JT, Barnette KG, Bohl CE, Hancock ML, Rodriguez D, Dodson ST, et al. The selective androgen receptor modulator GTx-024 (enobosarm) improves lean body mass and physical function in healthy elderly men and postmenopausal women: results of a double-blind, placebo-controlled phase II trial. J Cachexia Sarcopenia Muscle. 2011 Sep;2(3):153–61.

30. Wardley A, Cortes J, Provencher L, Miller K, Chien AJ, Rugo HS, et al. The ekicacy and safety of enzalutamide with trastuzumab in patients with HER2+ and androgen receptor-positive metastatic or locally advanced breast cancer. Breast Cancer Res Treat. 2021 May 1;187(1):155– 65.

31. Traina TA, Miller K, Yardley DA, Eakle J, Schwartzberg LS, O’Shaughnessy J, et al. Enzalutamide for the Treatment of Androgen Receptor–Expressing Triple-Negative Breast Cancer. JCO. 2018 Mar 20;36(9):884–90.

32. Yuan Y, Lee JS, Yost SE, Frankel PH, Ruel C, Egelston CA, et al. A Phase II Clinical Trial of Pembrolizumab and Enobosarm in Patients with Androgen Receptor-Positive Metastatic Triple-Negative Breast Cancer. The Oncologist. 2021 Feb 1;26(2):99–e217.

33. Yin D, Gao W, Kearbey JD, Xu H, Chung K, He Y, et al. Pharmacodynamics of Selective Androgen Receptor Modulators. J Pharmacol Exp Ther. 2003 Mar 1;304(3):1334–40.

34. Narayanan R, Ahn S, Cheney MD, Yepuru M, Miller DD, Steiner MS, et al. Selective Androgen Receptor Modulators (SARMs) Negatively Regulate Triple-Negative Breast Cancer Growth and Epithelial:Mesenchymal Stem Cell Signaling. PLOS ONE. 2014 Jul 29;9(7):e103202.

35. Yu Z, He S, Wang D, Patel HK, Miller CP, Brown JL, et al. Selective Androgen Receptor Modulator RAD140 Inhibits the Growth of Androgen/Estrogen Receptor–Positive Breast Cancer Models with a Distinct Mechanism of Action. Clinical Cancer Research. 2017 Dec 14;23(24):7608–20.

36. Hickey TE, Selth LA, Chia KM, Laven-Law G, Milioli HH, Roden D, et al. The androgen receptor is a tumor suppressor in estrogen receptor–positive breast cancer. Nat Med. 2021 Feb;27(2):310– 20.

37. Nyquist MD, Ang LS, Corella A, Coleman IM, Meers MP, Christiani AJ, et al. Selective androgen receptor modulators activate the canonical prostate cancer androgen receptor program and repress cancer growth. J Clin Invest. 131(10):e146777.

38. Mazzeo L, Ghosh S, Di Cicco E, Isma J, Tavernari D, Samarkina A, et al. ANKRD1 is a mesenchymal-specific driver of cancer-associated fibroblast activation bridging androgen receptor loss to AP-1 activation. Nat Commun. 2024 Feb 3;15(1):1038.

39. Korbecki J, Bosiacki M, Barczak K, Łagocka R, Brodowska A, Chlubek D, et al. Involvement in Tumorigenesis and Clinical Significance of CXCL1 in Reproductive Cancers: Breast Cancer, Cervical Cancer, Endometrial Cancer, Ovarian Cancer and Prostate Cancer. Int J Mol Sci. 2023 Apr 14;24(8):7262.

40. Ratna A, Das SK. Endothelin: Ominous Player in Breast Cancer. J Cancer Clin Trials. 2016 Feb;1(1):e102.

41. Lu JW, Liao CY, Yang WY, Lin YM, Jin SLC, Wang HD, et al. Overexpression of Endothelin 1 Triggers Hepatocarcinogenesis in Zebrafish and Promotes Cell Proliferation and Migration through the AKT Pathway. PLOS ONE. 2014 Oca;9(1):e85318.

42. Yin F, Bruemmer D, Blaschke F, Hsueh WA, Law RE, Herle AJV. Signaling pathways involved in induction of GADD45 gene expression and apoptosis by troglitazone in human MCF-7 breast carcinoma cells. Oncogene. 2004 Jun;23(26):4614–23.

43. Tadesse S, Yu M, Kumarasiri M, Le BT, Wang S. Targeting CDK6 in cancer: State of the art and new insights. Cell Cycle. 2015 Aug 28;14(20):3220–30.

44. Stucci LS, Internò V, Tucci M, Perrone M, Mannavola F, Palmirotta R, et al. The ATM Gene in Breast Cancer: Its Relevance in Clinical Practice. Genes (Basel). 2021 May 13;12(5):727.

45. Stagni V, Oropallo V, Barilà D. ATM: An unexpected tumor-promoting factor in HER2-expressing tumors. Mol Cell Oncol. 2015 Jun 10;3(2):e1054551.

46. Antonelli M, Strappazzon F, Arisi I, Brandi R, D’Onofrio M, Sambucci M, et al. ATM kinase sustains breast cancer stem-like cells by promoting ATG4C expression and autophagy. Oncotarget. 2017 Feb 20;8(13):21692–709.

47. Palomer X, Salvador JM, Griñán-Ferré C, Barroso E, Pallàs M, Vázquez-Carrera M. GADD45A: With or without you. Medicinal Research Reviews. 2024;44(4):1375–403.

48. Wang H, Guo M, Wei H, Chen Y. Targeting p53 pathways: mechanisms, structures and advances in therapy. Sig Transduct Target Ther. 2023 Mar 1;8(1):1–35.

49. Manousakis E, Miralles CM, Esquerda MG, Wright RHG. CDKN1A/p21 in Breast Cancer: Part of the Problem, or Part of the Solution? Int J Mol Sci. 2023 Dec 14;24(24):17488.

50. Lopez A, Reyna DE, Gitego N, Kopp F, Zhou H, Miranda-Roman MA, et al. Co-targeting of BAX and BCL-XL proteins broadly overcomes resistance to apoptosis in cancer. Nat Commun. 2022 Mar 7;13(1):1199.

51. Muluh TA, Shu X sheng, Ying Y. Targeting cancer metabolic vulnerabilities for advanced therapeutic ekicacy. Biomedicine & Pharmacotherapy. 2023 Jun 1;162:114658.

52. Sonkar K, Ayyappan V, Tressler CM, Adelaja O, Cai R, Cheng M, et al. Focus on the glycerophosphocholine pathway in choline phospholipid metabolism of cancer. NMR Biomed. 2019 Oct;32(10):e4112.

53. Barba I, Carrillo-Bosch L, Seoane J. Targeting the Warburg Ekect in Cancer: Where Do We Stand? International Journal of Molecular Sciences. 2024 Jan;25(6):3142.

54. Shin E, Koo JS. Glucose Metabolism and Glucose Transporters in Breast Cancer. Front Cell Dev Biol [Internet]. 2021 Sep 6 [cited 2025 Mar 9];9. Available from: https://www.frontiersin.org/journals/cell-and-developmental-biology/articles/10.3389/fcell.2021.728759/full

55. Khatami F, Payab M, Sarvari M, Gilany K, Larijani B, Arjmand B, et al. Oncometabolites as biomarkers in thyroid cancer: a systematic review. Cancer Manag Res. 2019 Feb 25;11:1829–41.

56. Azzman N, Anwar S, Syazani Mohamed WA, Ahemad N. Quinolone Derivatives as Anticancer Agents: Importance in Medicinal Chemistry. Curr Top Med Chem. 2024;24(13):1134–57.

57. Arabiyat S, Alzoubi A, Al-Daghistani H, Al-Hiari Y, Kasabri V, Alkhateeb R. Evaluation of Quinoline-Related Carboxylic Acid Derivatives as Prospective Dikerentially Antiproliferative, Antioxidative, and Anti-Inflammatory Agents. Chem Biol Drug Des. 2024 Oct;104(4):e14615.

58. Venugopala KN, Buccioni M. Current Understanding of the Role of Adenosine Receptors in Cancer. Molecules. 2024 Jan;29(15):3501.

